# Structural and binding studies of human CAMKK2 kinase domain bound to small molecule ligands

**DOI:** 10.1101/538157

**Authors:** Gerson S. Profeta, Caio V. dos Reis, Paulo H. C. Godoi, Angela M. Fala, Roger Sartori, Anita P. T. Salmazo, Priscila Z. Ramos, Katlin B. Massirer, Jonathan M. Elkins, David H. Drewry, Opher Gileadi, Rafael M. Couñago

## Abstract

Calcium/Calmodulin-dependent Protein Kinase Kinase 2 (CAMKK2) acts as a signaling hub, receiving signals from various regulatory pathways and decoding them via phosphorylation of downstream protein kinases - such as AMPK (AMP-activated protein kinase) and CAMK types I and IV. CAMKK2 relevance is highlighted by its constitutive activity being implicated in several human pathologies. However, at present, there are no specific small-molecule inhibitors available for this protein kinase. Moreover, CAMKK2 and its closest human homologue, CAMKK1, are thought to have overlapping biological roles. Here we present six novel co-structures of CAMKK2 bound to potent ligands identified from a library of ATP-competitive kinase inhibitors. Isothermal titration calorimetry (ITC) revealed that binding to some of these molecules is enthalpy driven. We expect our results to further advance current efforts to discover small molecule kinase inhibitors specific to each human CAMKK.

## Introduction

The Calcium/calmodulin-dependent kinases (CAMKs) respond to increases in the intracellular concentration of Ca^2+^ and play important roles in several cellular processes, mainly via the activation of transcription factors. These enzymes share a modular architecture composed of a kinase domain (KD) followed by an auto-inhibitory sequence and a partially overlapping calmodulin-binding domain (CBD). CAMKs are usually kept in an inactive state through auto-inhibition, which can be relieved, upon increases in Ca^2+^ levels, through the interaction of the protein CBD with Ca^2+^/calmodulin ^1^. CAMK family members include CAMK1-4 and two CAMK kinases (CAMKK1-2). As mediators of the second messenger effects of Ca^2+^, CAMK family members play prominent roles in cell division (CAMK3 aka eEF-2K - eukaryotic elongation factor 2 kinase); neuronal development (CAMK1, 2 and 4) and immune response (CAMK4) ^2,3^.

CAMK kinases (CAMKK) 1 and 2 are Ser/Thr kinases and upstream regulators of CAMKs ^4–6^. For example, CAMKKs phosphorylate and activate CAMK1 and CAMK4, resulting in the activation of cyclic AMP-responsive element binding protein (CREB). Consequently, both CAMKKs participate in several processes within the nervous system, such as neurite elongation and branching, long-term potentiation and memory ^7,8^.

In addition to regulating CAMK family members, CAMKK2 can phosphorylate AMP-activated protein kinase (AMPK). AMPK is a heterotrimeric protein complex that acts as an energy sensor and plays a key role regulating cellular energy metabolism. Dysregulation of AMPK has been implicated in major chronic diseases, such as obesity, inflammation, diabetes and cancer ^9^. AMPK activity can be allosterically controlled via competitive binding of different adenine nucleotides (ATP, ADP or AMP) to its regulatory gamma subunit ^10,11^. Alternatively, AMPK activation can be triggered by CAMKK2 via a nucleotide-independent mechanism. Thus, CAMKK2 can decode increases in intracellular Ca^2+^ levels triggered by upstream extracellular events, such as insulin receptor binding, to activate AMPK and maintain energy levels ^12–14^. Accordingly, CAMKK2-null mice are protected against weight gain induced by a high-fat diet, insulin resistance and glucose intolerance ^13^.

The development of potent and specific small molecule inhibitors to each of the CAMKKs would have a significant impact on our ability to investigate the roles these enzymes play in various biological processes. Due to the high degree of sequence and structure similarity between CAMKK1 and CAMKK2 ^15^, it is not surprising that most CAMKK inhibitors are equally potent against both enzymes ^16^. Moreover, STO-609 ^17^, a commonly used inhibitor of CAMKKs, demonstrates IC50 values less than 250 nM for six other kinases when screened against a panel of only 92 protein kinases ^18^. Thus, despite progress, the invention of a specific inhibitor for either CAMKK remains elusive.

Here we characterized the interaction of the kinase domain of CAMKK2 with several commercially available, small molecule kinase inhibitors. Our ITC data suggest that binding of the most potent compounds is enthalpy driven. We have also obtained co-crystal structures for CAMKK2 bound to some of these compounds. These co-structures suggest possible strategies for the development of CAMKK2-specific inhibitors.

## Results and Discussion

### DSF - Identification of kinase inhibitors that bind to CAMKK2

We employed a thermal-shift assay (Differential Scanning Fluorimetry, DSF)^19^ to identify compounds that can bind strongly to the kinase domain of CAMKK2. DSF screens were performed on purified CAMKK2 kinase domain (KD) (Supplementary Figure S1) using a library of commercially available, chemically diverse, ATP-competitive, kinase inhibitors. Compounds were available from two different laboratories (SGC-UNICAMP and SGC-Oxford) and DSF screens utilized two different formats (384- and 96-well, respectively). These differences in assay format prevented a direct comparison of absolute ∆Tm values found for compounds in each library. Nevertheless, compound ranking order obtained from either format is still informative. For example, structurally related compounds from different libraries (e.g., staurosporine and K252a) were found as top hits in both formats. Changes in melting temperature, ∆Tm, for CAMKK2-KD spanned a broad range of values (Fig. 1). Table 1 shows ∆Tm values for selected compounds (a comprehensive list can be found in Supplementary Table S1). Although not quantitative, DSF is a robust method to estimate binding affinities ^19^. Indeed, we found a good agreement between our DSF ∆Tm results and available *K*_D_ values from an unrelated competition binding assay using the full-length protein ^20^ (Supplementary Table S1). For example, staurosporine, a pan-kinase inhibitor, displayed the largest ∆Tm (17.1 °C) in our DSF assay and a *K*_D_ of 200 pM in the binding displacement assay. One notable discrepancy was Foretinib, a Type II kinase inhibitor originally developed to target members of the HGF and VEGFR receptor tyrosine kinase families ^21^. This compound displayed a high ∆Tm in our DSF assay (6.3 °C) and a low *K*_D_ (4,400 nM) in the binding displacement assay.

**Table 1.** CAMKK2 DSF, binding displacement and ITC data for selected compounds.

| Compound | $\Delta T_m$ (°C) <sup>a</sup> | ITC $K_D$ (nM) <sup>a</sup> | $K_D$ (nM) <sup>b</sup> |
| --- | --- | --- | --- |
| Staurosporine<sup>c</sup> | 17.1 | - | 0.2 |
| GSK650394<sup>c</sup> | 16.2 | 23.3 | - |
| ALK-IN-1<sup>c</sup> | 11.2 | 5.8 | - |
| Crenolanib (CP-868596) <sup>c</sup> | 10.6 | - | - |
| CP-673451<sup>c</sup> | 10.0 | - | - |
| TAE226<sup>c</sup> | 7.5 | - | - |
| Foretinib<sup>c</sup> | 6.3 | - | 4,400 |
| BI 2536<sup>c</sup> | 6.1 | 4.4 | 23 |
| BI 6727<sup>c</sup> | 6.0 | 8.1 | - |
| Crizotinib<sup>c</sup> | 3.3 | - | 1,500 |
| K252a<sup>d</sup> | 18.3 | - | - |
| TAE684<sup>d</sup> | 16.3 | - | 11 |
| Aminopurvalanol<sup>d</sup> | 9.7 | - | - |
| BIM I<sup>d</sup> | 7.2 | - | - |

a - this study; b - from Davies and colleagues, 2011; c - 384-well format; d - 96-well format

**Figure 1.**
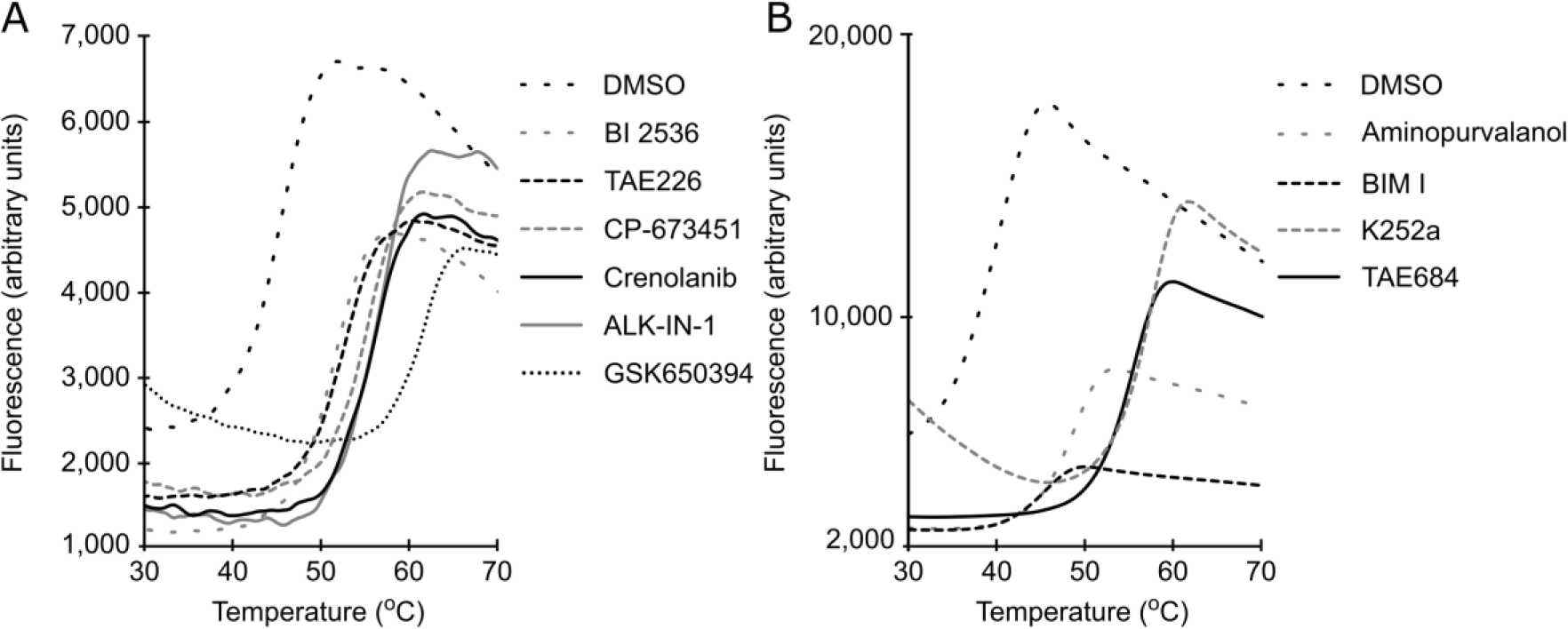
DSF screening identified ligands of CAMKK2-KD. Shown are CAMKK2-KD thermal melting curves in the absence (DMSO control) and presence of selected compounds from (**A**) 384-well and (**B**) 96-well format assays. ∆Tm values for these compounds can be found in Table 1 and Supplemental Table S1.

### Molecular basis for ligand binding

To better understand the ligand binding site of CAMKK2, we crystallized the protein kinase domain (amino acids 161-449) in the presence of some of our top DSF hits - BI 2536, ALK-IN-1, GSK650394, CP-673451, TAE226 and Crenolanib. All co-structures were solved by molecular replacement using CAMKK2-STO-609 ^22^ as search model. The resolution for these structures ranged from 1.8 to 2.0 Å (Table 2).

**Table 2.**
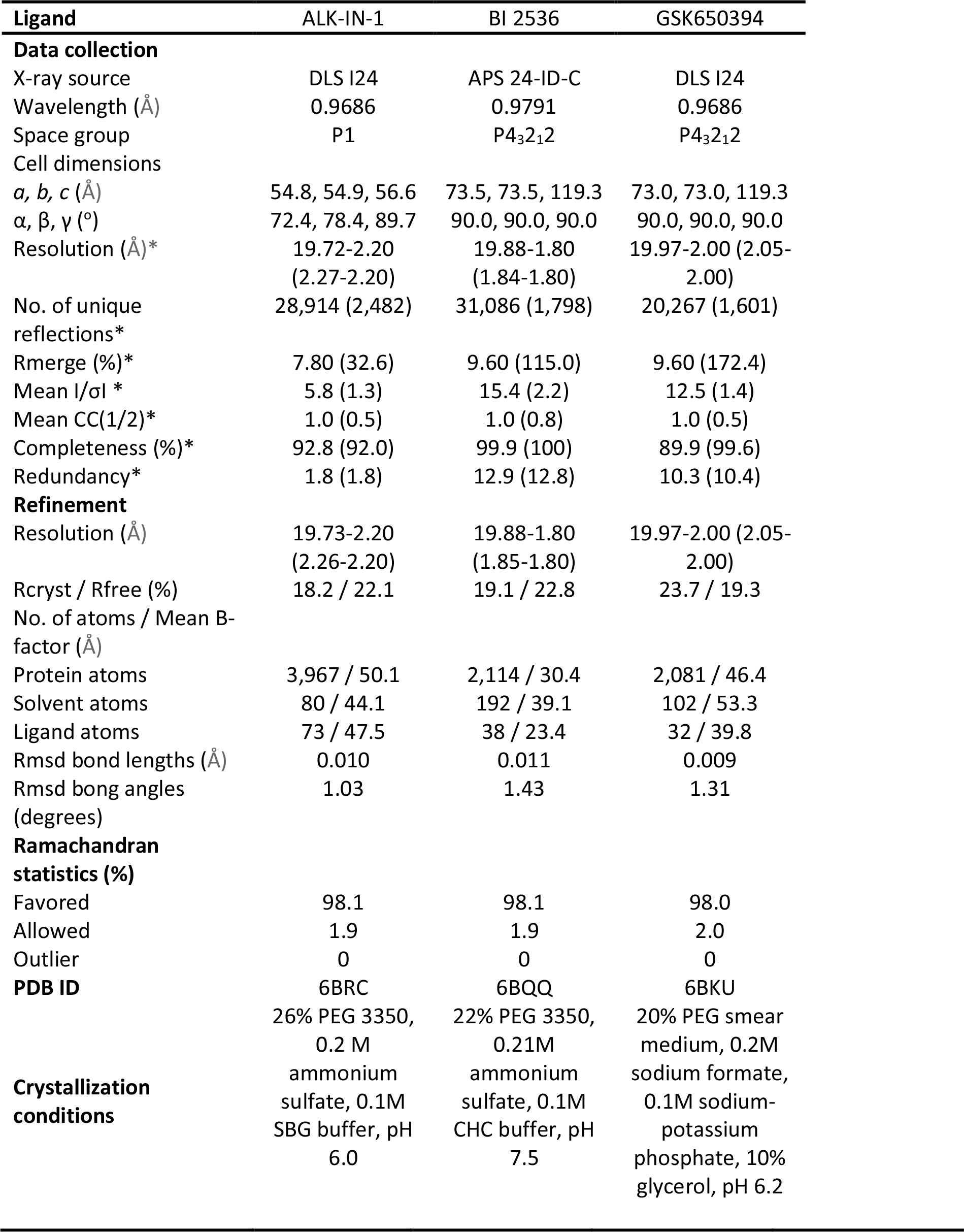
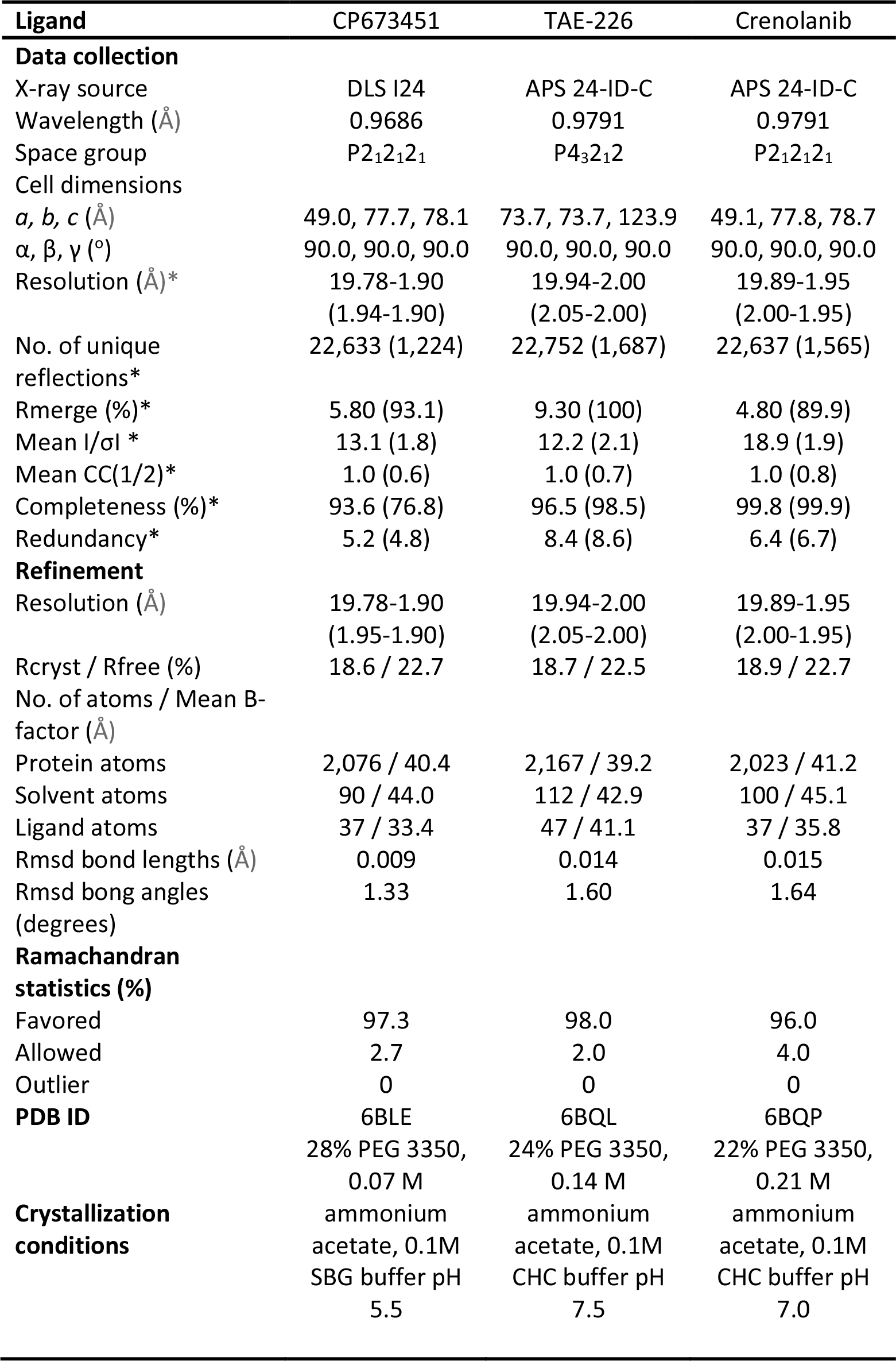
Crystallographic data.

For all co-structures the electron density maps were of good quality and allowed unambiguous building of the small molecule ligands within the protein ATP-binding site (Fig. 2). All compounds were anchored to the protein kinase hinge region via a hydrogen bond to the main-chain amide of residue Val270. For the majority of the compounds, an additional hydrogen bond was observed between the ligand and a carbonyl oxygen atom from the hinge region - Glu268 to GSK650394 or Val270 to TAE-226, ALK-IN-1 and BI 2536. When observed, all other polar interactions between protein and compound were water-mediated and located to the region between the ATP-binding site and the protein α-C.

**Figure 2.**
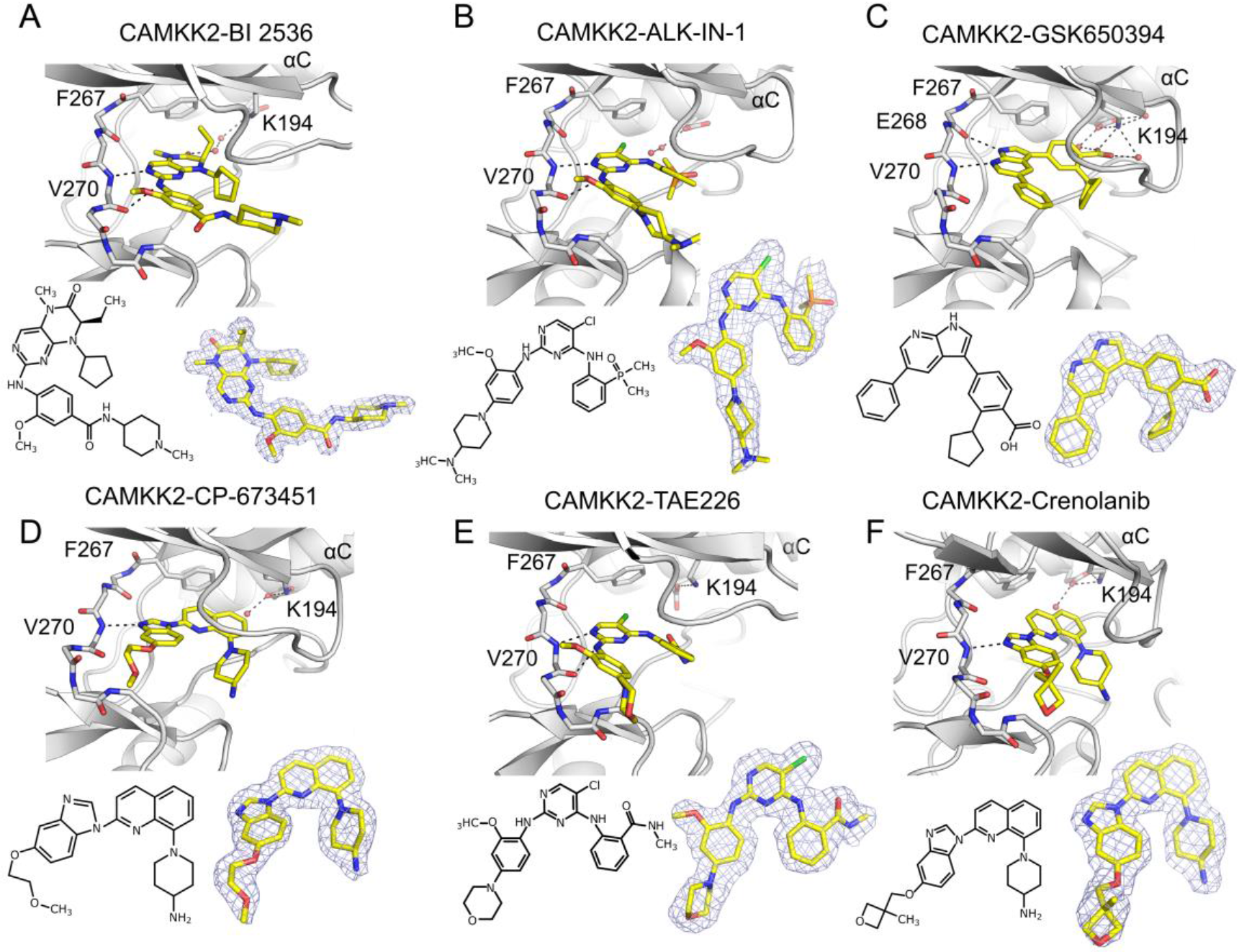
Binding mode of small-molecule ligands to CAMKK2-KD ATP-binding pocket. (**A-F**) Top part of each panel shows ligand binding mode and protein interactions. Main-chain atoms for residues (267-272) within CAMKK2-KD hinge region are represented as sticks. The side-chain atoms for CAMKK2-KD gatekeeper residue (Phe267) and catalytic lysine (Lys194) are also shown as sticks. Black dashed lines depict possible hydrogen bonds between protein and ligand atoms. Red spheres denote the position of crystallographic water molecules. The bottom part of each panel shows chemical structures and 2Fo-Fc electron density maps (contoured at 1.0 σ - gray meshes) for each ligand.

An overlay of all 6 co-structures revealed that the overall structure of CAMKK2-KD is mostly unchanged, despite being bound to different ligands. The most notable exception to this observation concerned the position of the protein P-loop. The P-loop is an inherently flexible (GxGxF/YG conserved motif) region of the kinase domain that folds over the phosphate atoms from the ATP molecule. For the different CAMKK2-KD co-structures obtained here, the distance between C-α atoms from P-loop residue Ser175 varied as much as 6.7 Å. We also observed a small change in the position of the α-C helix when bound to TAE-226 compared to the other co-structures (1.3 Å towards the ligand, Glu236 Cα-atom position) (Figure 3A). In all co-structures of CAMKK2-KD obtained here, the protein displayed an active (α-C-in) conformation, in which the R-spine is fully formed. The R-spine is a set of 4 residue in kinases whose hydrophobic side chains line up to form a spine when the enzyme is in an active conformation ^23^.

**Figure 3.**
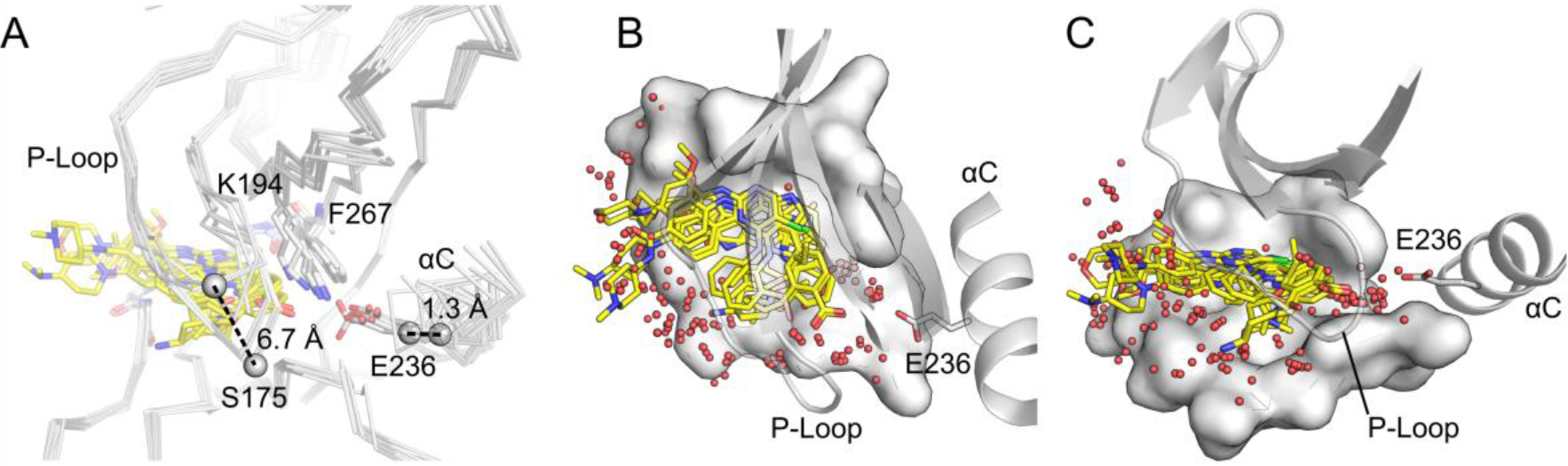
Structural Analysis of CAMKK2-KD bound to small-molecule kinase inhibitors. (**A**) Major structural differences between CAMKK2-KD bound to various inhibitors locate to the P-Loop and the α-C regions of the protein. Dashed lines indicate measurements between equivalent C-α atoms (distances are shown). (**B-C**) Top and side views of CAMKK2-KD ATP-binding site. Protein surface is shown for the bottom of the protein ATP-binding site. Shown is the CAMKK2-KD-GSK650394 structure. Ligands are depicted in stick model following the superposition of their cognate protein structures. Red spheres indicate solvent molecules from CAMKK2-KD structures reported in the literature (and bound to ALK-IN-1, BI 2536, GSK650394, CP673451, TAE226, Crenolanib, STO-609 and GSK650393 PDB IDs: 6BRC, 6BQQ, 6BKU, 6BLE, 6BQL, 6BQP, 2ZV2 and 6CMJ, respectively).

Despite the high resolution of these co-structures, we could not completely build the region between β-3 and α-C (residues 206-229). This region is known as the RP-insert and is rich in proline (8 out of 24), glycine (5 out of 24) and arginine (5 out of 24) residues, and likely to be disordered. The related human kinase, CAMMK1, also displays a similarly disordered region ^15^.

We compared our CAMKK2-KD co-crystal structures to those obtained for the same protein bound to STO-609 (PDB ID 2ZV2) ^22^ and a recently-developed azaindazole inhibitor similar to GSK650394 (PDB ID 6CMJ) ^16^. Overlay of all small molecules in the CAMKK2 ATP-binding site reveals that future compounds could explore the addition of polar groups to take place of the structural water molecules found trapped between the inhibitors and the conserved glutamate from the protein α-C helix (Figure 3B). Using inhibitor atoms to satisfy polar interactions originally facilitated between protein groups and crystallographic water molecules has been used successfully to increase compound potency in a number of cases. For example, Kung and colleagues used this strategy to increase the potency of compounds targeting HSP90 ^24^.

Nevertheless, potent inhibitors against both CAMKK proteins do exist ^16,17^ and a priority to better understand the function of these two proteins might be the development of specific inhibitors of each enzyme. Recently, our group obtained the structure of CAMKK1 and proposed structural differences between the 2 CAMKKs that could be exploited for the design of specific molecules to each protein. One of these strategies included taking advantage of the extra space observed for hinge residue Val270 in CAMKK2 ^15^. This suggestion that was also put forward by Price and colleagues ^16^.

### Favorable enthalpy dominates binding of most potent ligands to CAMKK2-KD

We performed ITC to confirm our DSF results and assess the thermodynamic parameters of ligand binding. We found that top hits on the DSF - BI 2536, ALK-IN-1, BI 6727 and GSK650394; were all potent binders of CAMKK2 and had *K*_D_ values ranging from 4.4 to 23.3 nM (Fig. 4). In general, *K*_D_ values from ITC experiments were in good agreement with our DSF ∆Tm values (Table 1). Nevertheless, GSK 650394 displayed the weakest binding by ITC (23.3 nM), but showed one of the largest ∆Tm values by DSF (16.2 °C). Further, an enzymatic IC_50_ of 600 pM was recently reported for compound GSK650394 (compound **1a** in Price and colleagues) against full-length CAMKK2 ^16^. Particular experimental conditions, such as the different buffers, enzymes and substrates employed in these experiments might be behind these differences.

**Figure 4.**
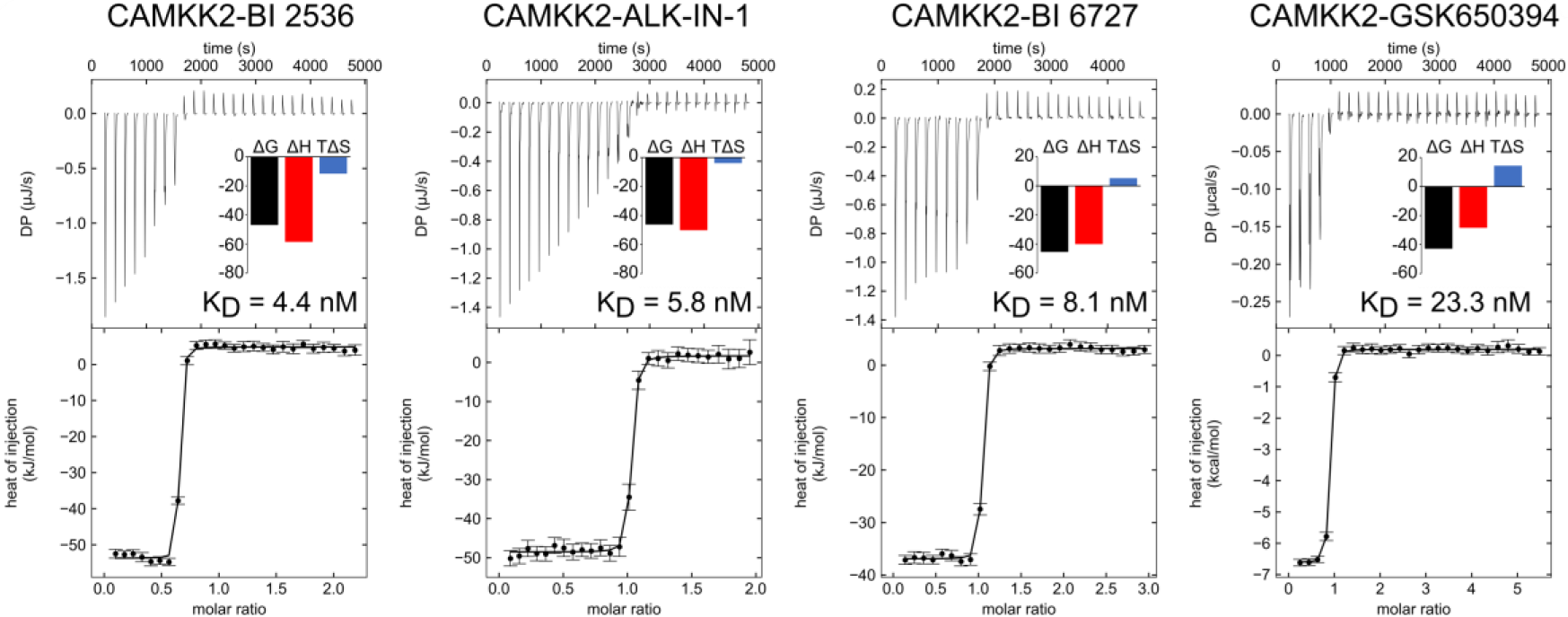
Dissociation constant measurements by ITC for selected compounds against CAMKK2-KD. The top part of each panel shows the injection heats, the bottom part shows the fitted binding isotherms (a single-site model was used), and the insets show the binding energies in kJ/mol.

Binding of all assayed compounds to CAMKK2-KD was enthalpy-driven, which is usually associated with the presence of hydrogen bonds and van der Waals interactions between ligand and protein ^25,26^. Indeed, all three compounds for which we obtained both ITC data and co-crystal structures (BI 2536, ALK-IN-1 and GSK650394) engaged the protein hinge region via two hydrogen bonds. These non-covalent interactions could account for the enthalpy values observed in our ITC experiments (ranging from −58.5 to −28.5 kJ/mol - Table 3). The fourth compound we obtained ITC data for, BI 6727, was originally described as a PLK1 inhibitor and interacts with its target enzyme via two hydrogen bonds to the protein hinge region ^27^. This compound is a close analogue of BI 2536, for which we were able to solve a structure. Assuming BI 6727 binding mode to CAMKK2-KD is the same as the one observed for PLK1 and comparable to our BI 2536 CAMKK2 structure, the enthalpy-driven interaction observed for this compound (−40.1 kJ/mol - Table 3) and CAMKK2-KD may also be explained by the formation of hydrogen bonds.

**Table 3.** Binding affinities and thermodynamic parameters measured by ITC.

|  |  |  |  |  |
| --- | --- | --- | --- | --- |
| **Compound** | **$K_D$**<br>**(nM)** | **$\Delta G$**<br>**(kJ/mol)** | **$\Delta H$**<br>**(kJ/mol)** | **$-T\Delta S$**<br>**(kJ/mol.K)** |
| GSK650394 | 23.3 | -42.8 | -28.5 | -14.4 |
| ALK-IN-1 | 5.8 | -46.2 | -50.1 | 3.9 |
| BI 2536 | 4.4 | -46.9 | -58.5 | 11.6 |
| BI 6727 | 8.1 | -45.4 | -40.1 | -5.4 |

### Conclusion

In conclusion we present here 6 novel co-structures of CAMKK2-KD bound to potent ligands selected from a library of ATP-competitive kinase inhibitors. Our co-structures showed that CAMKK2-KD had a similar binding mode with all ligands and point to possible design strategies for the invention of more potent and selective CAMKK2 inhibitors. An inhibitor specific for CAMKK2 would be able to set apart the biological functions of each one of the CAMKKs and further illuminate the different roles CAMKK2 plays in energy metabolism and in the nervous system.

## Methods

### Cloning, protein expression and purification

For crystallization of CAMKK2, we employed a construct of CAMKK2 isoform 7 residues 161-449 (NCBI NP_001257415.1 – SGC construct CAMKK2B-cb002) containing the wild-type kinase domain (CAMKK2-KD) in vector pNIC28-Bsa4. The construct was transformed into *Escherichia coli* BL21(DE3) cells that co-express λ-phosphatase and three rare tRNAs (plasmid pACYC-LIC+) ^28^. The cells were cultured in TB medium containing 50 μg/mL kanamycin and 34 μg/mL chloramphenicol at 37 °C with shaking until the OD_600_ reached ~3 and then cooled to 18°C for 1 hour. Isopropyl β-D-1-thiogalactopyranoside (IPTG) was added to a final concentration of 0.1 mM and the cultures were left overnight at 18°C. The cells were collected by centrifugation then resuspended in 2x lysis buffer [1x lysis buffer is 50 mM HEPES buffer, pH 7.5, 0.5 M KOAc, 10% (v/v) glycerol, 50 mM each arginine / glutamate, 10 mM imidazole, 1.0 mM tris(2-carboxyethyl)phosphine (TCEP), Protease Inhibitor Cocktail Set VII (Calbiochem, 1/500 dilution)] and flash-frozen in liquid nitrogen. For purification, the cell pellet was thawed and sonicated on ice. Polyethyleneimine (pH 7.5) was added to the lysate to a final concentration of 0.15% (w/v) and the sample was centrifuged at 53,000 × g for 45 min at 4 °C. The supernatant was loaded onto a Ni-Sepharose resin (GE Healthcare) and recombinant CAMKK2-KD was eluted stepwise in 1x lysis buffer with 300 mM imidazole. Removal of the hexahistidine tag was performed at 4°C overnight using recombinant TEV (Tobacco Etch Virus) protease. Protein was further purified using reverse affinity chromatography on Ni-Sepharose followed by gel filtration (Superdex 200 16/60, GE Healthcare). Protein in gel filtration buffer (10 mM HEPES, 500 mM KOAc, 1.0 mM TCEP, 5% [v/v] glycerol) was concentrated to 9 mg/mL (measured by UV absorbance in a NanoDrop spectrophotometer (Thermo Scientific) using the calculated molecular weight and estimated extinction coefficient) using 30 kDa molecular weight cut-off centrifugal concentrators (Millipore) at 4°C. Concentrated protein was flash-frozen in a liquid nitrogen bath and stored at −80 °C until use.

### Crystallization, data collection and structure determination

Kinase inhibitors (dissolved in 100 % DMSO) were added to the protein in a 3-fold molar excess and incubated on ice for approximately 30 minutes. The mixture was centrifuged at 21,000 x g for 10 minutes at 4°C prior to setting up 150 nL volume sitting drops at three ratios (2:1, 1:1, or 1:2 protein-inhibitor complex to reservoir solution). Crystallization experiments were performed at 20 °C. Crystals were cryoprotected in mother liquor supplemented with 25-30% glycerol before flash-cooling in liquid nitrogen for data collection. Diffraction data was collected at 100 K at the Advanced Photon Source 24ID-C, and at the Diamond Light Source beam line I24. Data collection statistics and crystallization conditions can be found in Table 2.

Diffraction data was integrated with XDS ^29^ and scaled using AIMLESS from the CCP4 software suite ^30^. The structure was solved by molecular replacement using Phaser ^31^ and the kinase domain of CAMKK2 as the search model (PDB ID 2ZV2) ^22^. Refinement was performed using REFMAC5 ^32^ and Coot ^33^ was used for model building. Structure validation was performed using MolProbity ^34^.

### Differential scanning fluorimetry (DSF)

*384-well format* - CAMKK2-KD protein was screened against a library of 378 structurally diverse ATP-competitive kinase inhibitors available from Selleckchem (Houston, TX, United States; catalog No. L1200). Each well contained 20 μL of 1 μM kinase in 100 mM potassium phosphate pH 7.0, 150 mM NaCl, 10% glycerol and the Applied Biosystems Protein Thermal Shift dye at the recommended concentration of 1:1000.

The compounds, previously solubilized in DMSO, were used at 10 μM final concentration and 0.1% DMSO. Plates were sealed using optically clear films and transferred to a QuantStudio 6 qPCR instrument (Applied Biosystems). The fluorescence intensity was measured during a temperature gradient from 25 to 95 °C at a constant rate of 0.05 °C/s, and protein melting temperatures were calculated based on a Boltzmann function fitting to experimental data, as implemented in the Protein Thermal Shift Software (Applied Biosystems). Protein in 0.1% DMSO was used as a reference.

*96-well format* - Each well contained 20 μL of 2 μM kinase in 10 mM HEPES pH 7.5 and 500 mM NaCl. SYPRO Orange dye (Invitrogen) was used in 5X as final concentration.

The compounds, previously solubilized in DMSO, were used at 12 μM final concentration and 2.5% DMSO. Plates were sealed using optically clear films and transferred to an Agilent Mx3005p RT-PCR machine (Agilent). The fluorescence intensity was measured during a temperature gradient from 25 to 95 °C at a constant rate of 0.05 °C/s, and protein melting temperatures were calculated based on a Boltzmann function fitting to experimental data (Graphpad Prism 7). Protein in 2.5% DMSO was used as a reference.

For both formats, compounds that caused a shift in melting temperature of the protein (∆Tm) of 2 °C or higher compared to the reference were considered positive hits.

### Isothermal titration calorimetry

Measurements were made using a MicroCal AutoITC200 (Malvern, United Kingdom), at 20 °C with 1,000 rpm stirring. For all measurements CAMKK2-KD protein was dialyzed overnight against gel filtration buffer, and the dialysis buffer was used to dilute the inhibitors. CAMKK2-KD protein was titrated into a solution containing the inhibitor. The concentrations used for each measurement were: BI 2536 and BI 6727 at 24.3 μM; and CAMKK2-KD at 243.6 μM; and ALK-IN-1 and GSK650394 at 20.1 μM, and CAMKK2-KD at 201 μM. All measurements were made using 1.5 μL injections and 180 s between each injection. ITC data was analyzed with NITPIC, and SEDPHAT; figures were made using GUSSI ^35^.

## Acknowledgements

The SGC is a registered charity (number 1097737) that receives funds from AbbVie, Bayer Pharma AG, Boehringer Ingelheim, Canada Foundation for Innovation, Eshelman Institute for Innovation, Genome Canada, Innovative Medicines Initiative (EU/EFPIA) [ULTRA-DD grant no. 115766], Janssen, Merck KGaA Darmstadt Germany, MSD, Novartis Pharma AG, Ontario Ministry of Economic Development and Innovation, Pfizer, Takeda, and Wellcome [106169/ZZ14/Z]. This work was supported by the Brazilian agencies FAPESP (Fundação de Amparo à Pesquisa do Estado de São Paulo) (2013/50724-5 and 2014/5087-0) and CNPq (Conselho Nacional de Desenvolvimento Científico e Tecnológico) (465651/2014-3). This work was funded in part by the National Cancer Institute of the National Institutes of Health (grant number R01CA218442). This work is based upon research conducted at the Northeastern Collaborative Access Team beamlines (GU51510, GU56413), which are funded by the National Institute of General Medical Sciences from the National Institutes of Health (P41 GM103403). The Pilatus 6M detector on 24-ID-C beam line is funded by a NIH-ORIP HEI grant (S10 RR029205). This research used resources of the Advanced Photon Source, a U.S. Department of Energy (DOE) Office of Science User Facility operated for the DOE Office of Science by Argonne National Laboratory under Contract No. DE-AC02-06CH11357. We thank Diamond Light Source for access to beamlines I-24 (MX16171) that contributed to the results presented here. We thank the staff of the Life Sciences Core Facility (LaCTAD) from State University of Campinas (UNICAMP), for the Genomics and Proteomics analysis. PZR, AMF, and CVR received CAPES (Coordenação de Aperfeiçoamento de Pessoal de Nível Superior) fellowships (88887.136432/2017-00, 88887.136437/2017-00 and 88887.146077/2017-00, respectively). RS and GSP received FAPESP technical training fellowships (2016/09041-0 and 2017/05697-0, respectively).

## Author Contributions

G.S.P performed protein purification, crystallization and ITC experiments. R.S. performed crystallization experiments. R.M.C. performed protein purification and crystallization, and collected diffraction data. C.V.R. and R.M.C solved co-crystal structures. A.P.T.S., P.Z.R. and O.G. designed constructs and did cloning. A.M.F., P.H.C.G. and C.V.R. performed compound screening and binding data analysis. K.B.M., O.G., D.H.D., J.M.E. and R.M.C. coordinated the project. R.M.C. wrote the manuscript. All authors revised the manuscript.

## Corresponding author

Correspondence to Rafael M. Couñago.

## Data Availability Statement

The coordinates and structure factors for all CAMKK2-KD co-crystal structures reported here have been deposited in the Protein Data Bank with accession codes 6BRC, 6BQQ, 6BKU, 6BLE, 6BQL and 6BQP.

---

**Supplementary Figure S1.**
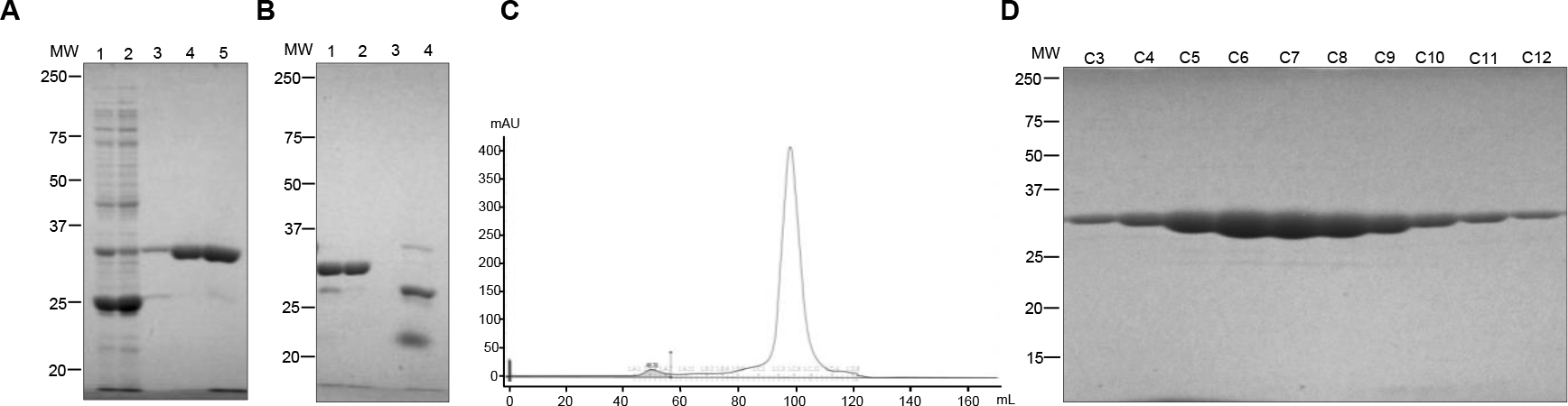
Results for a typical CAMKK2-KD purification. **A**-Electrophoretic analysis of IMAC fractions: soluble fraction (lane 1), Ni-NTA flow-through fraction (lane 2), Ni-NTA wash in 10 mM imidazole (lane 3), Ni-NTA wash in 30 mM imidazole (lane 4) and eluate with 300 mM imidazole from Ni2+ (lane 5). **B**-SDS-PAGE analysis of Ni-NTA eluted fractions after TEV protease treatment (lane 1). Ni unbound protein Flow-Through (lane 2), Ni-NTA wash in 30 mM imidazole (lane 3) and Ni-NTA eluate in 300 mM imidazole (lane 4). **C**-Gel filtration chromatogram. **D**-SDS-PAGE analysis of gel filtration samples. Precision Plus Protein Unstained Protein Standards (Bio-Rad, cat no. 161-0363) was used as molecular weight marker.

## Supplementary Information

**Table S1.**
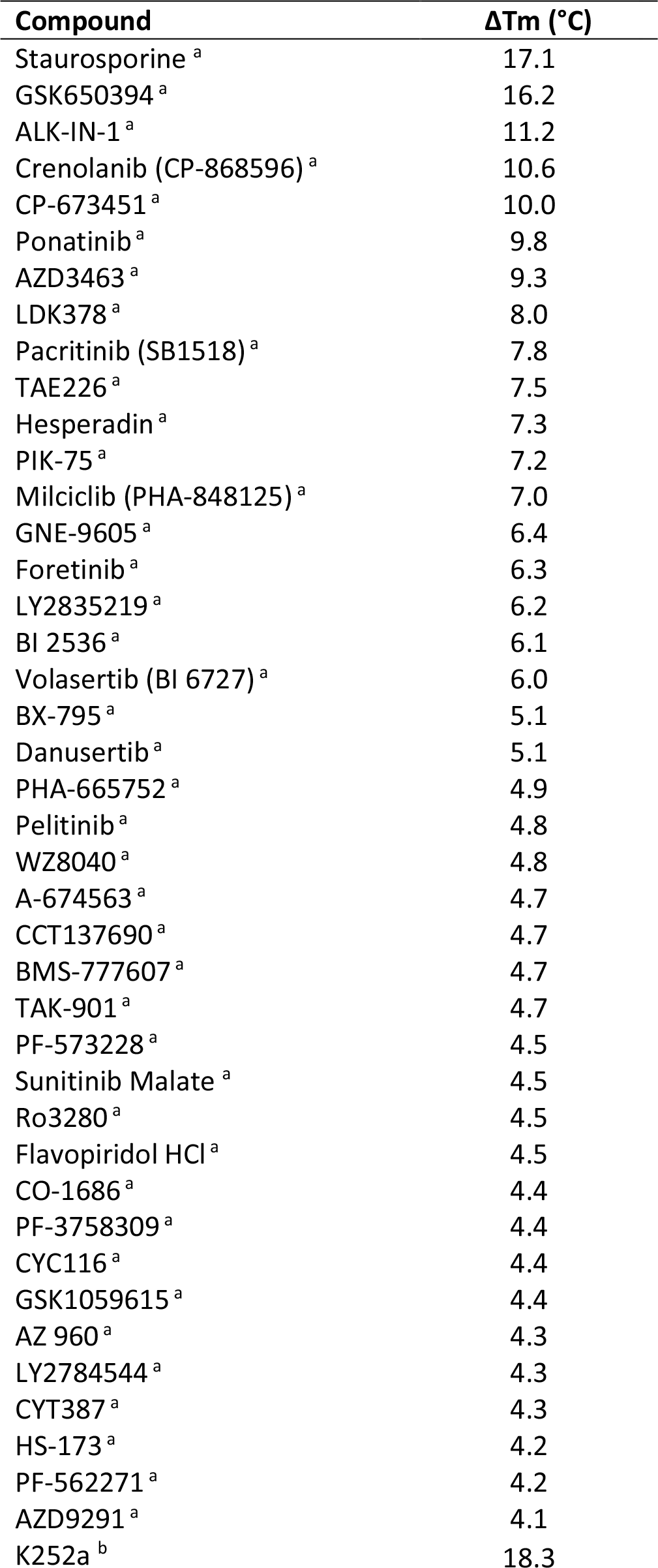
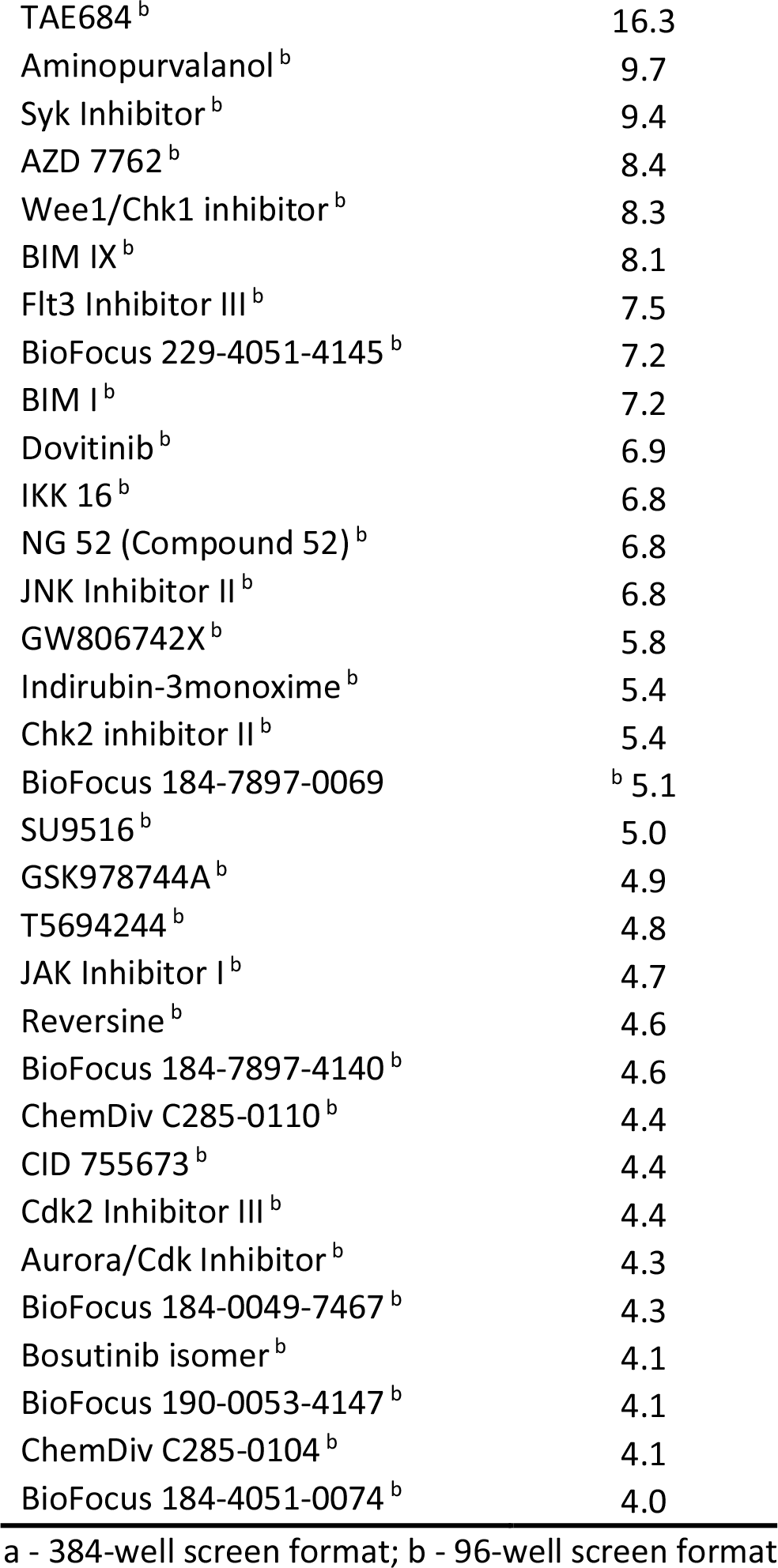
DSF screening results for compounds that increased the Tm of CAMKK2 > 4.0 °C.

